# Granulosa cell-layer stiffening prevents the granulosa cells from escaping the post-ovulatory follicle

**DOI:** 10.1101/2024.02.06.579251

**Authors:** Xiaodong Wang, Jianning Liao, Hongru Shi, Yongheng Zhao, Wenkai Ke, Hao Wu, Guoshi Liu, Xiang Li, Changjiu He

## Abstract

Ovulation is necessary for successful reproduction. After ovulation, cumulus cells and oocytes are released, while granulosa cells (GCs) remain trapped within the post-ovulatory follicle to form the corpus luteum. However, the mechanism underlying GC confinement has long been unclear. Here, we provide *in vitro* and *in vivo* evidence demonstrating that the stiffening of GC-layer as an evolutionarily conserved mechanism that hinders GCs from escaping the post-ovulatory follicles. Spatial transcriptome analysis reveals that the assembly of focal adhesions is primarily responsible for this stiffening. Disrupting focal adhesion assembly through RNA interference results in the release of GCs from the post-ovulatory follicle, leading to the formation of an aberrant corpus luteum with reduced cell density and cavities. We also uncover that the *LH (hCG) -cAMP-PKA-CREB* signaling axis stimulates focal adhesion assembly and induce GC-layer stiffening. Our findings introduce a novel concept of “GC-layer stiffening”, which offers valuable insights into the factors that prevent GCs escape from the post-ovulatory follicle.

## INTRODUCTION

Ovulation, a fundamental event in female reproduction, signifies not only the culmination of oogenesis but also the commencement of luteinization. This event is triggered by ovulatory signals, specifically the luteinizing hormone (LH) surge or human chorionic gonadotropin (hCG). The pre-ovulatory follicle, which has the potential to ovulate, is a sophisticated structure consisting of GCs, cumulus cells, oocyte, and theca cells. Each cell type plays a unique role in coordinating the programmed ovulation [1, 2].

Upon receiving an ovulatory signal, the follicular cell types undergo distinct fates. The oocyte resumes meiosis, becomes fertile, and is released from the ruptured follicle [3, 4]. In parallel, the cumulus cell-layer undergoes extracellular matrix (ECM) remodeling, which leads to its expansion and increased viscosity [5]. In addition, the cumulus cells also produce inflammatory mediators and chemokines, creating an inflammatory microenvironment that aids in follicle rupture [6, 7]. Eventually, cumulus cells accompany the oocyte during its escape to the fallopian tube’s ampulla. In contrast, GCs primarily function in receiving and transmitting the ovulatory signal. Abundant LH receptors on the GC cytomembrane allow for sensitive recognition of the signal [8, 9]. Moreover, signaling cascades like the “EGF-like factor signaling pathway” and “MAPK3/1 signaling pathway” in GCs play a crucial role in amplifying the ovulatory signal and transmitting it to cumulus cells and oocyte. However, unlike cumulus cells and oocytes, GCs cannot escape the follicle and instead remain within the post-ovulatory follicle to form the corpus luteum, which regulates the estrus cycle and is essential for maintaining pregnancy.

Interestingly, during folliculogenesis, cumulus cells and GCs originate from a common progenitor in preantral follicles [12, 13]. It is intriguing, therefore, to consider why cumulus cells are able to escape from the post-ovulatory follicle, while GCs, with a shared cellular origin, are unable to do so. This question is quite puzzling, and thus far, no theoretical model has been developed to explain it.

This study aims to establish a theoretical model to address this question. We discovered that the GC-layer undergoes a process called “GC-layer stiffening” upon receiving the signal. This stiffening prevents the GC-layer from escaping the ruptured follicle. Through spatial transcriptome sequencing and conducting *in vitro* and *in vivo* experiments, we confirmed that the assembly of focal adhesions, triggered by the *LH (hCG)-cAMP-PKA-CREB* signaling cascade, is crucial for “GC-layer stiffening”. Disrupting focal adhesion assembly through RNA interference led to a failure in “GC-layer stiffening” and subsequent release of GCs from the post-ovulatory follicle. This resulted in the formation of an abnormal corpus luteum with low cell density and cavitation.

## RESULTS

### 1. Ovulatory signal triggered the stiffening of GC-layer and inhibited its escape from the punctured follicle

To facilitate real-time studying and monitoring of the ovulation process, we developed a mouse follicle culture system capable of supporting ovulation and luteinization while allowing for gene knockdown within the follicle (Figure 1A). By puncturing the cultured follicles before or after the addition of hCG, we observed distinct outcomes in the release of GCs. Puncturing before hCG addition or 1 hour after hCG addition resulted in easy release of GCs, while at 6 and 10 hours post hCG addition, minimal or no GCs were released (Figure 1B, Movie 1). Importantly, we observed a significant increase in the rigidity of the GC-layer following hCG addition. The GC-layers exhibited low rigidity at H0 and H1, fragmenting under mechanical oscillation, but demonstrated enhanced rigidity at H6 and H10, enabling them to maintain structural integrity when exposed to the same concussive force (Figure 1C). We termed this phenomenon “GC-layer stiffening”.

**Figure 1.**
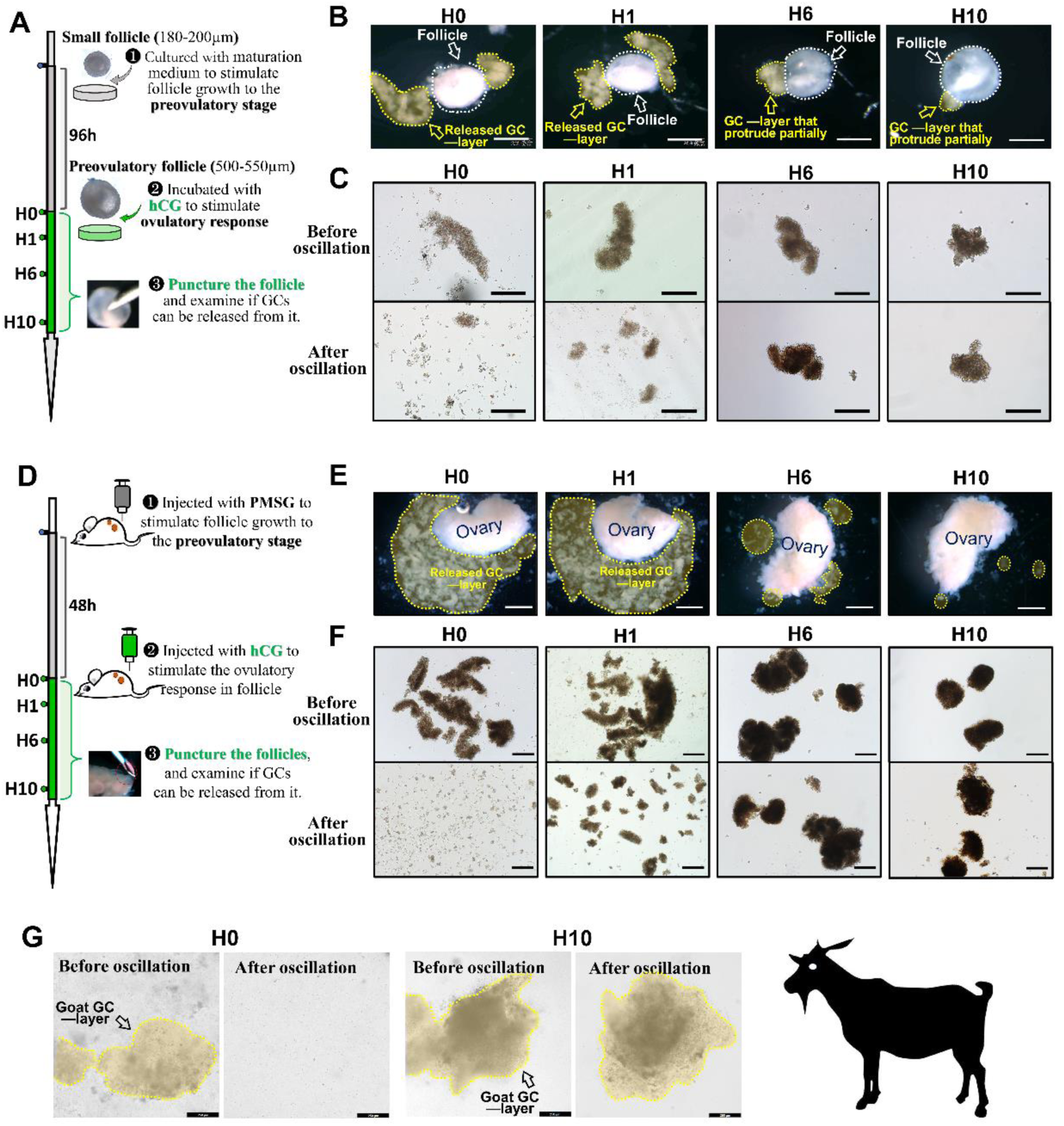
Ovulatory signal triggered the stiffening of GC-layer and inhibited its escape from the punctured follicle. A, Experimental design of *B* and *C*. B, Effect of hCG addition on the capability of GC-layer to escape from punctured follicles. Scale bar: 400 µm. GC-layers outlined by yellow frames, and follicles outlined by white frames. C, Effect of hCG addition on the rigidity of GC-layer. Oscillation parameter: 700rpm, 1 min, and 37 ℃. Scale bar: 400 µm. D, Experimental design of *E, F*. E, Effect of hCG injection on the capability of GC-layer to escape from punctured ovaries. Scale bar: 1mm. GC-layers outlined by yellow frames. F, Effect of hCG injection on the rigidity of GC-layer. Oscillation parameter: 700 rpm, 1 min, and 37 ℃. Scale bar: 400 µm. G, Effect of hCG injection on the rigidity of goat GC-layer. Oscillation parameter: 700rpm, 1 min, and 37 ℃. Scale bar: 250 µm. GC-layers outlined by yellow frames. B, C, E, F were repeated independently five times, and G was repeated two times. Similar results were observed.

We further investigated “GC-layer stiffening” *in vivo* using the superovulation technique (Figure 1D). Consistent with our *in vitro* findings, hCG injection induced “GC-layer stiffening” and prevented GC release from the punctured ovary. Briefly, at H0 and H1, the unstiffened GC-layer burst out of the punctured ovarian surface (Figure 1E; Movie 2) and disintegrated upon mechanical oscillation (Figure 1F, Movie 3), while at H6 and H10, the stiffened GC-layer remained trapped in the ovary with only a few GCs released (Figure 1E, Movie 2). These released GCs remained intact after mechanical oscillation (Figure 1F, Movie 3). Our observations led to the hypothesis that the ovulatory signal-triggered “GC-layer stiffening” determines GC escape from the follicle.

Interestingly, we also observed “GC-layer stiffening” in goats. Before hCG injection, the GC-layer in goats exhibited low rigidity, disintegrating after mechanical oscillation. However, at 10 hours post hCG injection, the GC-layer displayed increased rigidity and remained intact (Figure 1G).

### 2. Spatial transcriptome analysis suggested that focal adhesion could potentially serve as the structural foundation for GC-layer stiffening

Considering the diverse cell types and follicles at different developmental stages within the ovary, we conducted spatial transcriptomic analysis of the ovaries of H0 and H6 to explore the mechanisms underlying the occurrence of “GC-layer stiffening”. Spatial transcriptome data was obtained from public databases [14]. Following rigorous quality control procedures and personalized data analysis, we identified 7 pre-ovulatory follicles at H0 and 10 at H6, respectively. From these follicles, we obtained transcriptional changes of GCs following hCG stimulation (Figure 2A). Principal component analysis (PCA) demonstrated significant transcriptional differences among these GCs (Figure 2B). A total of 5352 differentially expressed genes were identified using the DESeq2 software. Among these genes, 2303 were found to be up-regulated after hCG injection (Figure 2C). Gene ontology (GO) analysis demonstrated that the up-regulated genes were mainly enriched in biological processes related to cell connection, such as *Focal Adhesion*, *Cell-cell Junction*, *Integrin Binding*, and *Anchoring Junction*, in addition to well-known ovulation-related processes (Figure 2D). This led us to hypothesize that alterations in cell connection may contribute to “GC-layer stiffening”.

**Figure 2.**
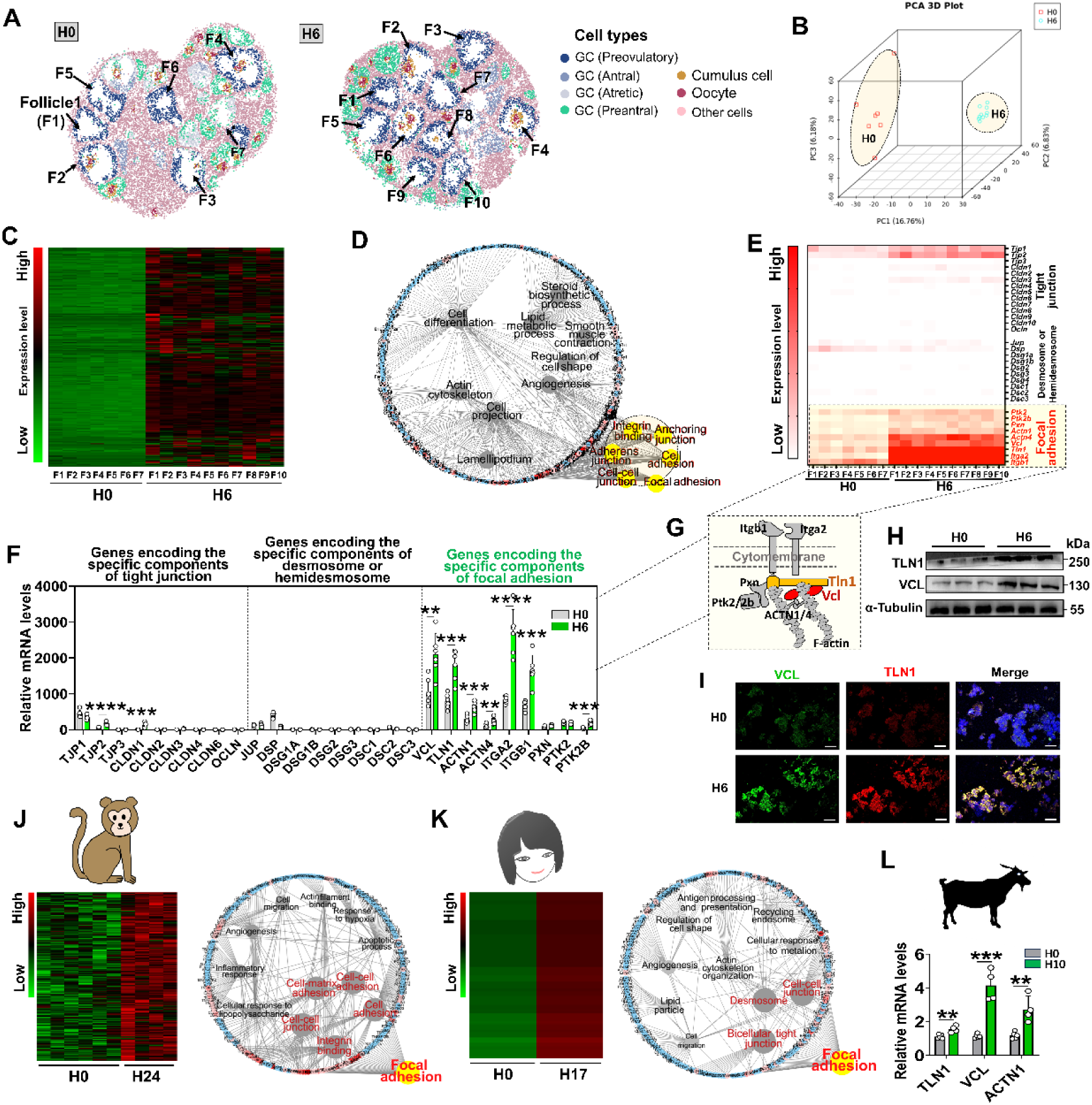
Spatial transcriptome analysis suggested that focal adhesion could potentially serve as the structural foundation for GC-layer stiffening. A, Identification of pre-ovulatory follicles within the ovaries through spatial transcriptome analysis. n = 7 (H0) and 10 (H6) pre-ovulatory follicles, respectively. The GCs within the pre-ovulatory are highlighted in blue. B, PCA analysis of the transcriptome discrepancy in GCs within pre-ovulatory follicles. C, Heat map of the up-regulated genes in GCs after hCG injection. D, GO analysis of the up-regulated genes. Biological processes related to cell connection outlined by yellow frames. E, Heat map of genes encoding the components of tight junction, desmosome, hemidesmosome and focal adhesion. F, qRT-PCR validation of the expression of genes encoding the components of tight junction, desmosome, hemidesmosome, and focal adhesion. n = 6 GC samples. G, Schematic representation of the structure of focal adhesion. H, Western blot assay of the protein contents of VCL and TLN1 after hCG injection. n = 3 GC samples. Original blots can be viewed in Fig. S4A. I, Immunofluorescence analysis of the localization of VCL and TLN1 in GC-layer after hCG injection. Scale bar: 20 µm. J, Analysis of the transcriptome changes in monkey GC after hCG injection. *Left*: Heat map of the up-regulated genes; *right*: GO analysis of the up-regulated genes. Focal adhesion outlined by yellow. K, Analysis of the transcriptome changes in human GC after hCG injection. *Left*: Heat map of the up-regulated genes; *right*: GO analysis of the up-regulated genes. Focal adhesion outlined by yellow. L, qRT-PCR analysis of the expression of goat genes encoding the components of focal adhesion after hCG injection. n = 5 (H0) and 4 (H10) GC samples. Statistical significance was determined using two-tailed unpaired Student’s t test, values were mean ±SD. **P<0.01, ***P<0.001, ****P<0.0001. F, H and I were repeated independently three times, similar results were obtained.

To investigate the specific type of cell connection involved in achieving “GC-layer stiffening,” we examined genes encoding components of tight junction, desmosome, hemidesmosome, and focal adhesion. Analysis of the spatial transcriptomic data showed that genes encoding components of focal adhesions were highly expressed and upregulated by hCG in GCs (Figure 2E). To validate the transcriptome data, we performed quantitative real-time PCR (qRT-PCR), which exhibited consistent gene expression patterns (Figure 2F). Moreover, Western blotting and immunofluorescence confirmed the significant induction of VCL and TLN1 (Figure 2H, I), core structural proteins of focal adhesion (Figure 2G), in the GC-layer after hCG injection. Analysis of transcriptome data from other species revealed that focal adhesion assembly during ovulation is not exclusive to mice but also occurs in monkey and human (Figure 2J, K). qRT-PCR assay also showed upregulation of *VCL*, *TLN1*, and *ACTN1* in goat GCs after hCG injection (Figure 2L). These findings indicate that ovulatory signal-induced focal adhesion assembly is a conserved event across species. We hypothesized that the ovulatory signaling induces focal adhesion assembly, leading to the occurrence of “GC-layer stiffening”.

### 3. Disruption of focal adhesion assembly led to a failure of the stiffening of GC-layer and an escape of GCs from the manually punctured follicle

To substantiate our hypothesis and elucidate the functional role of focal adhesion assembly in GC-layer stiffening, we employed lentivirus-mediated RNA interference to knock down the expression of *VC*L and *TLN1* in cultured follicles (Figure 3A). The result indicated that transfecting plasmids 48 hours prior to the addition of ovulation-inducing medium can effectively silence the expression of *VCL* and *TLN1* at H6 (Figure S1), without exerting adverce impact on the growth of follicles to the preovulatory stage (Figure 3B). However, the knockdown of *VCL* and *TLN1*, either individually or in combination, led to the unstable retention of GCs within the follicles of H6. Upon puncturing the follicles, GCs in the *si-VCL* (Movie 4), *si-TLN1* (Movie 5), and *si-VCL+TLN1* (Movie 6) groups easily burst out, while those in the control groups struggled to escape (Figure 3C). Furthermore, we discovered that the disruption of focal adhesion assembly hindered the stiffening of the GC-layer, as the GC-layers in the *si-VCL*, *si-TLN1*, and *si-VCL+TLN1* groups showed lower rigidity and disintegrated upon mechanical oscillation compared to the control groups (Figure 3D).

**Figure 3.**
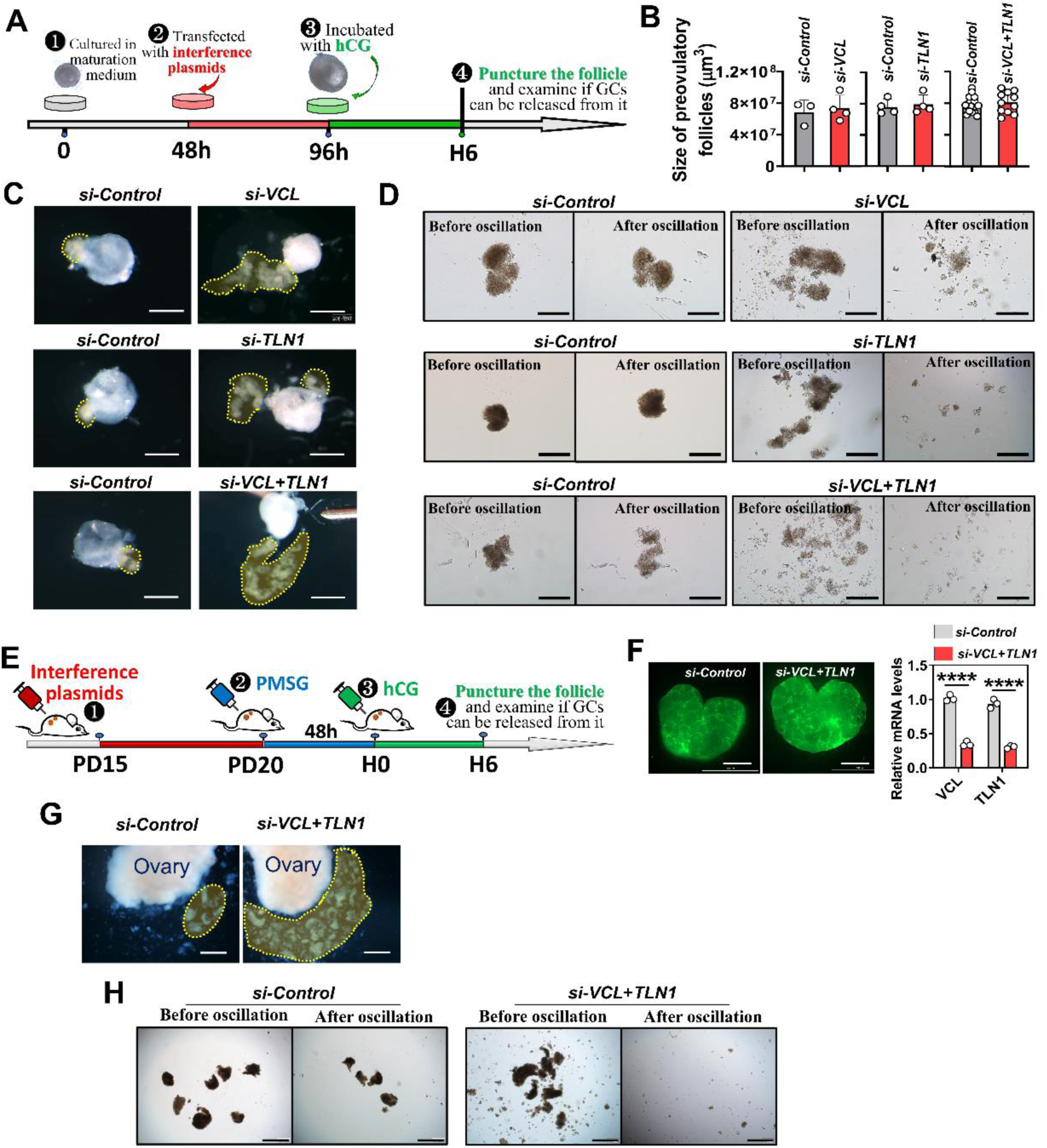
Disruption of focal adhesion assembly led to a failure of the stiffening of GC-layer and an escape of GCs from the punctured follicle. A, Schematic representation of the knockdown of *VCL* and *TLN1* in cultured follicles. B, Effect of *VCL* and *TLN1* knockdown on follicle growth, n = 3 (*si-Control*), 4 (*si-VCL*); 4 (*si-Control*), 4 (*si-TLN1*); 18 (*si-Control*), 11 (*si-VCL+TLN1*). The scrambled shRNA was used as *si-Control* in this study. C, Effect of *VCL* and *TLN1* knockdown on the capability of GC-layer to escape from the punctured follicles. Scale bar: 400 µm. GC-layers outlined by yellow frames. D, Effect of *VCL* and *TLN1* knockdown on the rigidity of GC-layer. Oscillation parameter: 700 rpm, 1 min, and 37 ℃. Scale bar: 400 µm. E, Schematic representation of the knockdown of *VCL* and *TLN1* in ovaries. F, qRT-PCR analysis of the efficiency of *VCL*+*TLN1* interference. Green fluorescence indicates successful transcription of interfering plasmids in ovaries. Scale bar: 1 mm, n=3 ovaries, collected from 3 mice. G, Effect of *VCL*+*TLN1* knockdown on the capability of GC-layer to escape from punctured ovaries. Scale bar: 1 mm. GC-layers outlined by yellow frames. H, Effect of *VCL*+*TLN1* knockdown on the rigidity of GC-layer. Oscillation parameter: 700 rpm, 1 min, and 37 ℃. Scale bar: 800µm. Statistical significance was determined using two-tailed unpaired Student’s t test. Values were mean ± SD. ****P<0.0001. C, D were repeated independently five times, and G, H repeated two times. Similar results were observed.

To further validate our findings, we performed injections of lentiviral particles beneath the ovarian bursa to specifically knock down the expression of *VCL* and *TLN1* in the ovaries (Figure 3E, F). Consistent with our *in vitro* observations, simultaneous knockdown of *VCL* and *TLN1* facilitated the release of GC-layers from the ovaries at H6 (Figure 3G). Notably, the GC-layer in the *si-VCL+TLN1* group exhibited reduced rigidity, leading to its disintegration upon mechanical oscillation compared to the control group (Figure 3H). These findings provide compelling evidence supporting the pivotal role of focal adhesion assembly, induced by the ovulatory signal, in the stiffening of the GC-layer and its subsequent inability to escape from manually punctured follicles.

### 4. Disruption of focal adhesion assembly resulted in the release of GCs from the post-ovulatory follicle and a reduction in the quantity of luteal cells

Figure 3 showed that the disruption of focal adhesion assembly led to the failure of “GC-layer stiffening” and resulted in the release of GCs from punctured follicles. However, it is imperative to ascertain whether the disruption of focal adhesion assembly leads to the spontaneous release of GCs from the post-ovulatory follicle, akin to the cumulus-oocyte complex (COC). To address this, real-time filming of ovulation was conducted (Figure 4A). We observed that the simultaneous knockdown of *VCL* and *TLN1* had no significant on ovulation rate (Figure 4B). However, compared to the control group where only the COC was expelled (Figure 4C/left, outlined by green frames), the GC-layer in the *si-VCL+TLN1* group prominently protruded from the rupture site during COC expulsion (Figure 4C/left, outlined by yellow frames, Movie 7). Remarkably, following complete COC expulsion, a substantial number of free cells flowed out from the rupture site (Figure 4C/left, outlined by red frames, Movie 7), which were identified as GCs through qRT-PCR analysis of marker gene *Lhcgr* (Figure 4C/right). Subsequently, we evaluated the morphology and function of the newly formed corpus luteum at H40. Compared to the control group, the knockdown groups (*si-VCL*, *si-TLN1*, and *si-VCL+TLN1*) exhibited lower cell density and distinct cavitation in the corpus luteum (Figure 4D, S2). Additionally, the expression levels of luteal functional genes and progesterone content were significantly reduced in the knockdown groups compared to the control groups (Figure 4E, S2). These findings provide direct evidence supporting the pivotal role of focal adhesion-mediated “GC-layer stiffening” in confining GCs within the post-ovulatory follicle for the purpose of differentiating into a corpus luteum.

**Figure 4.**
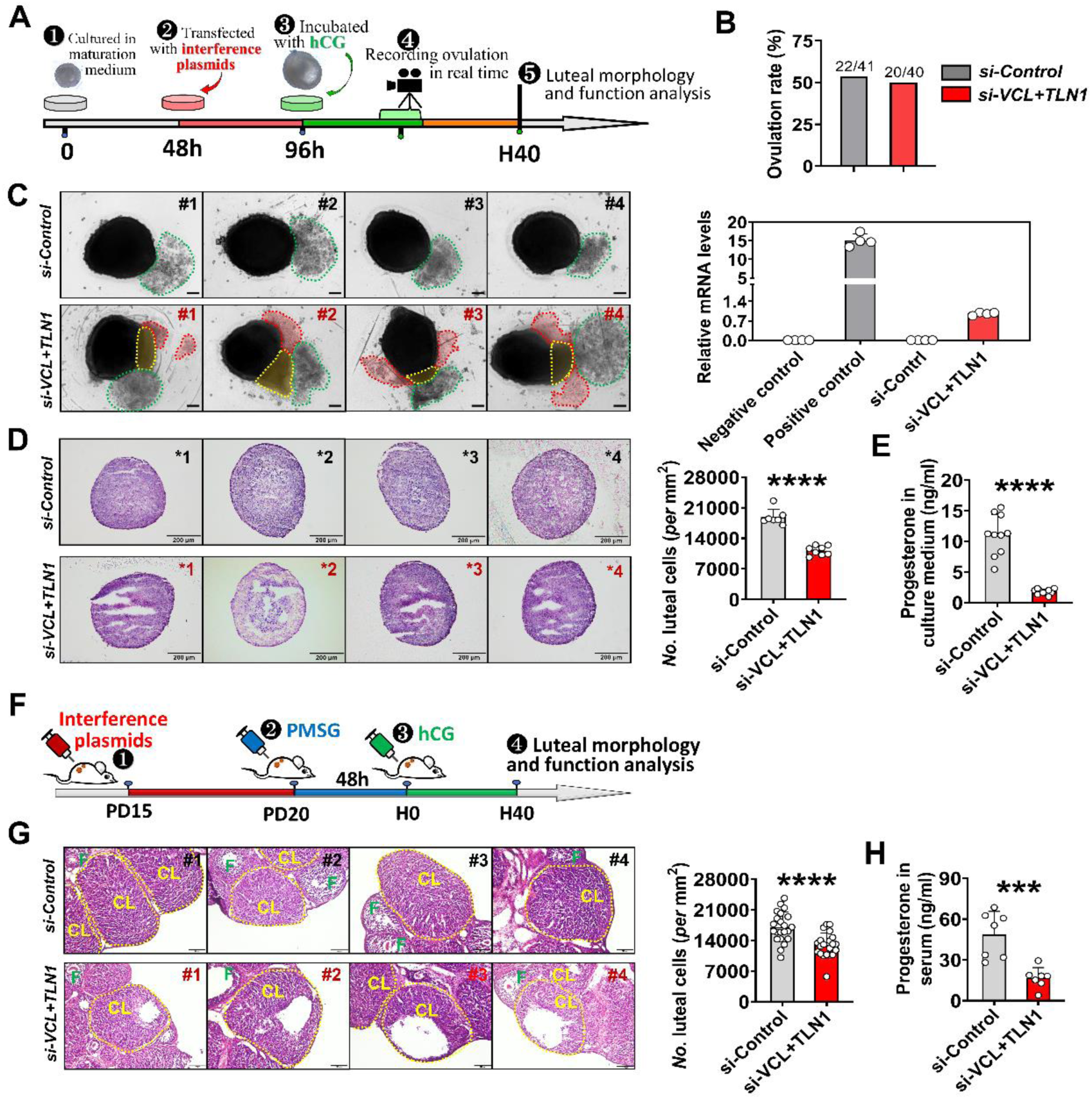
Disruption of focal adhesion assembly resulted in the release of GCs from the post-ovulatory follicle and a reduction in the quantity of luteal cells. A, Schematic representation for real-time recording of ovulation process after *VCL+TLN1* knockdown. B, Effect of *VCL+TLN1* knockdown on the ovulation rate. C, Knocking down *VCL+TLN1* results in the spontaneous release of GCs from the post-ovulatory follicle. *Left*: Representative photographs of the post-ovulatory follicles. Scale bar: 100 µm. The free GCs released from the rupture site are outlined by red frames. GC clumps protruded from the rupture site are outlined by yellow frames. The released COCs are covered by green frames. *Right*: identity verification of the released GCs through qRT-PCR. n=4 GC samples. *Lhcgr* was chosen as the marker gene for GC. Purified GCs and cumulus cells were used as positive and negative controls, respectively. D, Effect of *VCL+TLN1* knockdown on the morphology and function of the *in vitro* corpus luteum. *Left*: representative photographs of luteal sections in each group. Scale bar: 200 µm. *Right*: statistics of the density of luteal cells in each group, n = 7 (*si-Control*), 8 (*si-VCL+TLN1*). E, Effect of *VCL+TLN1* knockdown on progesterone level in culture medium. n = 10 medium samples in each group. F, Experimental design of *G*, *H*. G, Effect of *VCL+TLN1* knockdown on the morphology and function of the *in vivo* corpus luteum. *Left*: representative photographs of ovarian sections. CL = corpus luteum, which are outlined by yellow frames. F = follicle. Scale bar: 100 µm. *Right*: statistics of the density of luteal cells, n = 22 (*si-Control*), 21 (*si-VCL+TLN1*) CL. These CLs was observed from 4 and 7 biological independently ovaries, respectively. H, Effect of *VCL+TLN1* knockdown on progesterone level in serum. n = 7 serum samples in each group. Statistical significance was determined using two-tailed unpaired Student’s t test and Chi-squared test, values were mean ± SD. ***P<0.001, ****P<0.0001. C and D was repeated independently five times, G was repeated two times. Similar results were obtained.

We also validated our findings through an *in vivo* knockdown experiment. Simultaneous knockdown of *VCL* and *TLN1* in the ovaries (Figure 4F) resulted in a notable decrease in the cell density of the newly formed corpus luteum, accompanied by the presence of cavities within these corpus lutea (Figure 4G). Additionally, the average serum progesterone content in the *si-VCL+TLN1* mice was only 30% of that in the control mice (Figure 4H). These abnormal phenotypes are consistent with those observed *in vitro*.

### 5. Ovulatory signal stimulated focal adhesion assembly and “GC-layer stiffening” by activating *cAMP-PKA-CREB* cascade

To determine the signaling pathways involved in focal adhesion assembly and “GC-layer stiffening”, we analyzed genes upregulated by hCG (Figure 2C) using Kyoto Encyclopedia of Genes and Genomes (KEGG) analysis. This analysis revealed multiple signaling pathways, with the top three being the “*MAPK*”, “*cAMP*”, and “*PI3K-AKT*” (Figure 5A). In parallel, we used JASPAR (http://jaspar.genereg.net/) to predict the transcription factors binding to the promoters of focal adhesion structural genes in mice. Interestingly, CREB, a core transcription factor in *cAMP-PKA* signaling pathway, showed high binding scores (Figure S3A). Moreover, by analyzing ChIP-seq data of CREB derived from human embryonic stem cells (http://cistrome.org/db/#/, GSM1010896), we identified significant binding peaks of CREB in the promoters of six focal adhesion structural genes, including *VCL*, *TLN1*, *ACTN1*, *ACTN4*, *ITGA2* and *ITGB1* (Figure 5B). Based on these observations, we hypothesized that the *cAMP-PKA-CREB* signaling cascade may play a pivotal role in stimulating focal adhesion assembly and “GC-layer stiffening”.

**Figure 5.**
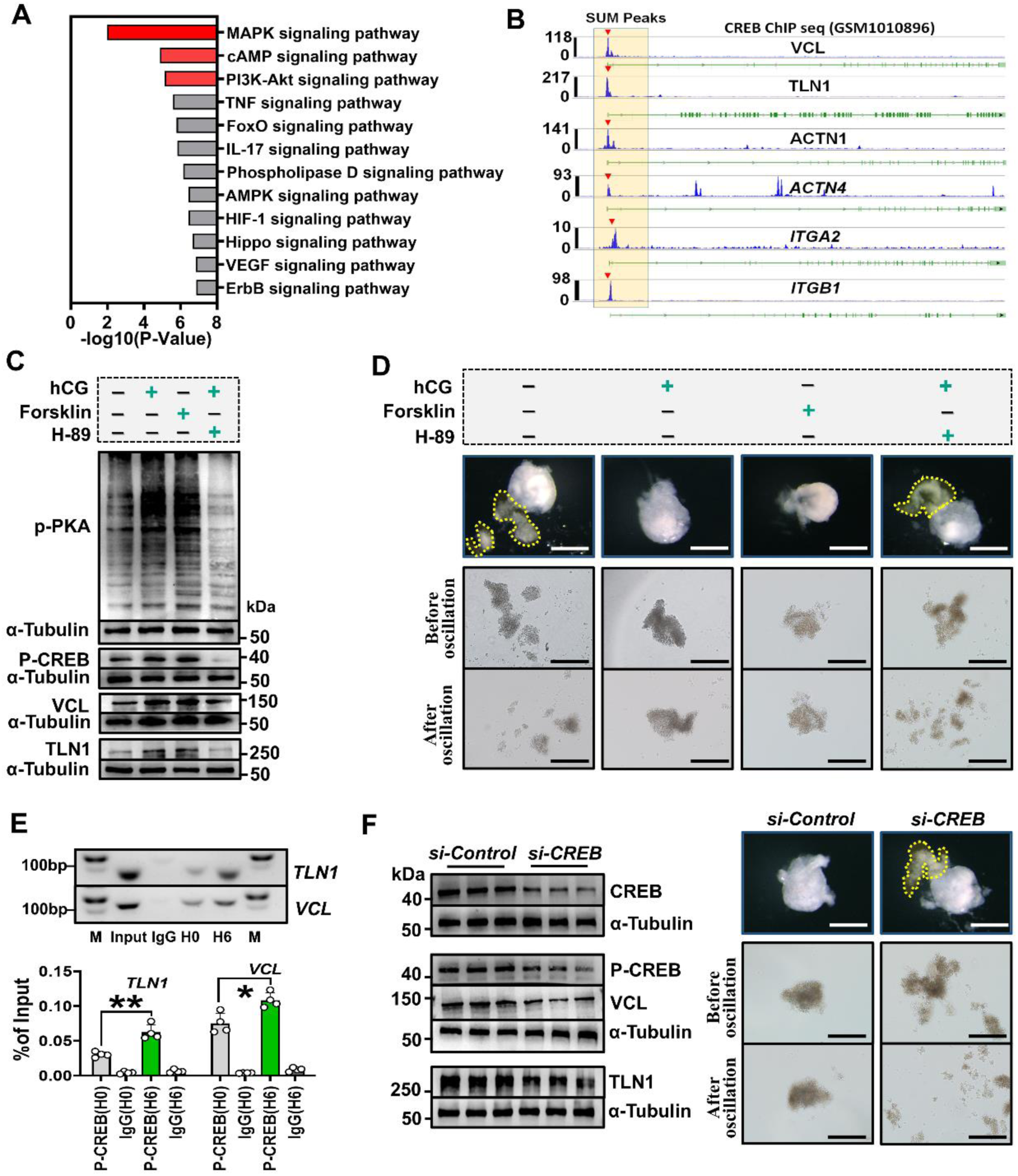
Ovulatory signal stimulated focal adhesion assembly and “GC-layer stiffening” by activating *cAMP-PKA-CREB* cascade. A, KEGG analysis of the up-regulated genes. B, Analysis of the binding of CREB to promoter of focal adhesion structural genes. The binding peaks of CREB are indicated by red trangles. C, Western blot assay of the protein contents of VCL and TLN1 following activation or inhibition of the *cAMP-PKA* cascade. Original blots can be viewed in Fig.S4B. D, Effect of activating or inhibiting *cAMP-PKA* cascade on the rigidity and escape capability of the GC-layer. Scale bar: 400 µm. The released GC-layers are outlined by yellow frames. E, ChIP-qPCR assay for CREB binding to the promoters of *VCL* and *TLN1*. *Up*: electrophoretic images of PCR products. Input and IgG were used as positive and negative controls, respectively. Original gel images can be viewed in Fig.S5. *Down*: statistical chart of qPCR assay. n = 4 GC samples. F, Effect of *CREB* knockdown on focal adhesion assembly and “GC-layer stiffening”. *Left*: western blot assay of protein contents of VCL and TLN1 after *CREB* knockdown. Original blots can be viewed in Fig.S4C. *Right*: Effect of *CREB* knockdown on the rigidity and escape capability of the GC-layer. Scale bar: 400 µm. The released GC-layers are outlined by yellow frames. Statistical significance was one-way ANOVA followed by Tukey’s post hoc test, values were mean ± SD. *P<0.05, **P<0.01. C, D and F were repeated independently three times, E was repeated two times. Similar results were observed.

To test this hypothesis, we conducted experiments using our follicle culture system. At H4, we observed that forskolin, an activator of adenylate cyclase, was sufficient to increase VCL and TLN1 protein levels by activating *PKA-CREB*, even without hCG addition (Figure 5C). This activation resulted in “GC-layer stiffening” and prevented the escape of GCs from punctured follicles at H6 (Figure 5D). Conversely, the use of H89, a PKA inhibitor, effectively suppressed *PKA-CREB* activity, preventing hCG-induced increases in VCL and TLN1 levels (Figure 5C) and subsequent “GC-layer stiffening” (Figure 5D). Furthermore, through ChIP-qPCR and dual-luciferase reporter assays, we determined the specific motifs that directly bind to CREB in the promoters of *VCL* and *TLN1* as CCAGGATGGCCTCAAACTTT and CAAGAGTGACATCATACACT, respectively. Notably, the bindings of CREB to these motifs significantly increased 6 hours after hCG addition (Figure 5E, Figure S3B, C). Lastly, we performed a knockdown of *CREB* expression in cultured follicles. Compared to the control group, the knockdown of *CREB* resulted in a significant reduction in VCL and TLN1 levels at H6, and more importantly, it led to the failure of “GC-layer stiffening” and subsequently the release of GCs from punctured follicles (Figure 5F). Altogether, these results strongly confirm our speculation that the *LH (hCG)-cAMP-PKA-CREB* signaling pathway is a key regulator of focal adhesion assembly and “GC-layer stiffening”.

## DISCUSSION

GCs and cumulus cells, despite arising from the same progenitor cells, have distinct fates after ovulation. While cumulus cells are released along with the oocyte, GCs must remain within the follicle to transform into the corpus luteum. The mechanism preventing GC escape from the post-ovulatory follicle has long been unclear. In this study, we propose a concept called “GC-layer stiffening” to explain this phenomenon (Figure 6). “GC-layer stiffening” is triggered by the ovulation signal, specifically the *LH (hCG)-cAMP-PKA-CREB* pathway. Prior to the ovulation signal, the GC-layer is in a flexible state, allowing for easy escape. However, upon ovulation signal stimulation, the GC-layer undergoes stiffening, preventing escape. Blockade of “GC-layer stiffening” results in the release of GC release from the post-ovulatory follicle and the formation of an abnormal corpus luteum characterized by low cell density and cavitation. Remarkably, we observed “GC-layer stiffening” in goats as well (Figure 1G), indicating its evolutionary conservation among higher mammals.

**Figure 6.**
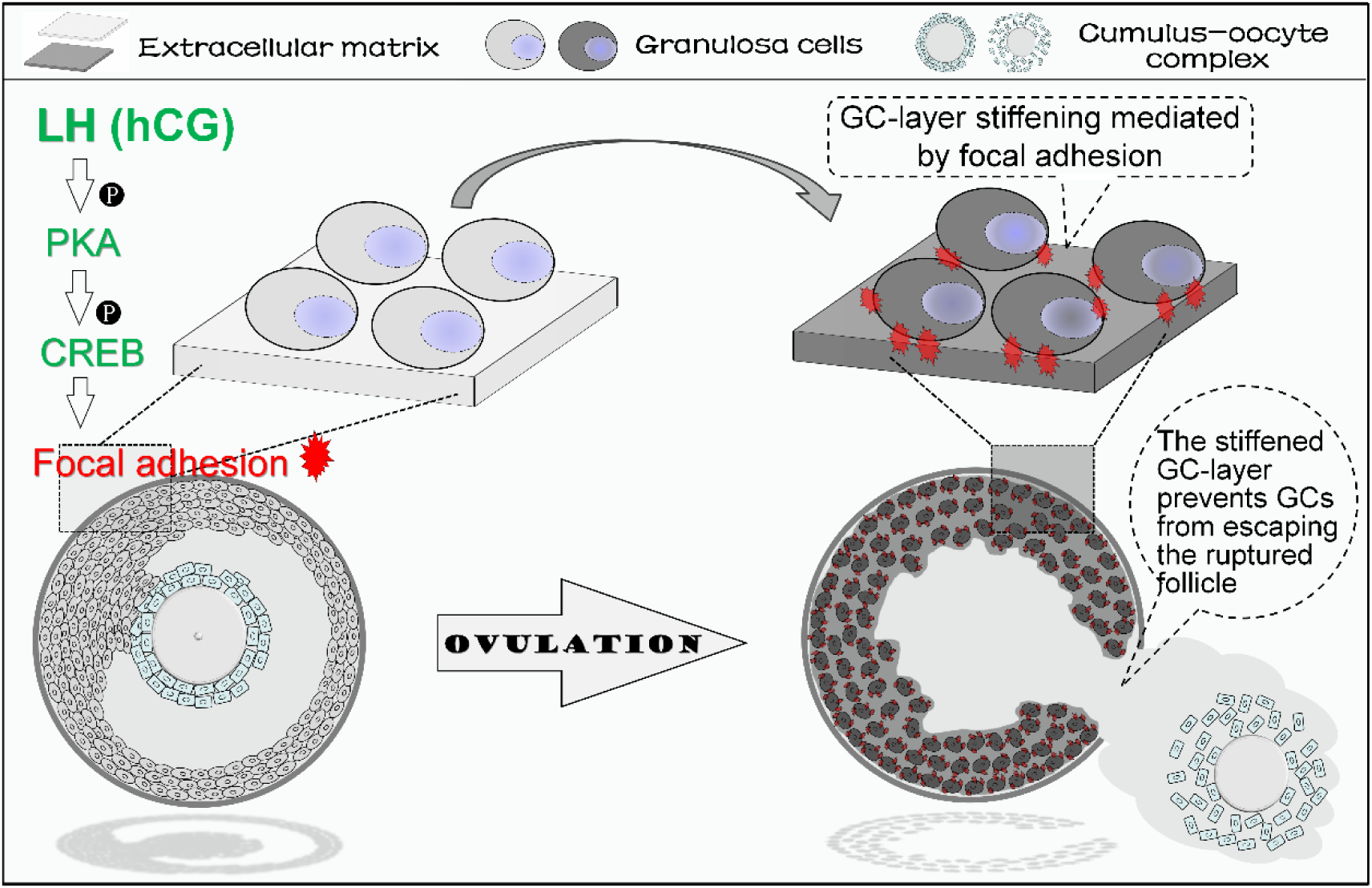
A diagram depicting “GC-layer stiffening” preventing GCs from escaping a post-ovulatory. LH or hCG stimulation enhances the *cAMP-PKA-CREB* signaling pathway in GCs of the pre-ovulatory follicle. This pathway triggers the assembly of focal adhesions within the GC layer, leading to a significant increase in the strength of the connections between GCs and the extracellular matrix (ECM). As a result, the GC layer becomes more rigid and stiff. This increased rigidity reduces the flexibility of the GC layer, preventing it from escaping through the ruptured follicle.

Why does stiffening have the capacity to prevent GCs from escaping the post-ovulatory follicle? We propose that the increased rigidity it imparts to the GC-layer is key. During ovulation, contractile activity and ECM proteolysis facilitate the release of the cumulus-oocyte complex (COC) from the rupture site [15]. However, we observed in Movie 7 that the rupture site is smaller than the size of the COC, resulting in deformation upon exit. The ability of the COC to pass through the narrow rupture site relies on its flexibility, which is achieved through cumulus expansion. In contrast, the stiffened GC-layer lacks flexibility, hindering its ability to exit through the narrow opening. Previous studies have shown that ECM stiffness plays a role in regulating cell behaviors through mechanotransduction mechanisms [16-18]. However, the specific role of “GC-layer stiffening” in activating signaling networks within the pre-ovulatory follicle was not explored in this study. Notably, inhibition of “GC-layer stiffening” resulted in a decrease in the expression of functional genes in lutein cells derived from residual GCs (Figure S2). This suggests that, beyond preventing GC escape, blocking “GC-layer stiffening” may disrupt the signaling network involved in luteinization. Therefore, it is important to investigate the downstream signaling pathways triggered by “GC-layer stiffening” in the pre-ovulatory follicle.

We consider that the assembly of focal adhesions serves as a structural foundation for “GC-layer stiffening”, as evidenced by the failure in “GC-layer stiffening” when focal adhesion assembly is disrupted (Figure 3). Remarkably, we also observed ovulatory signal-triggered focal adhesion assembly in goat, monkey, and human specimens (Figure 2J-L), indicating that focal adhesion-mediated “GC-layer stiffening” is a conserved mechanism. It should be noted that focal adhesions also play a vital role in cell migration [19], thereby the observed cavitation of the corpus luteum in the *si-VCL+TLN1* group (Figure 4) may be attributed to a combination of GC escape and impaired migration. While investigating other adhesion structures that may contribute to “GC-layer stiffening”, we found extremely low expression levels of the structural genes of desmosome and hemidesmosome, known for their involvement in cell-cell and cell-ECM adhesion, respectively [20, 21], in GCs (Figure 2), ruling out their significant involvement. However, we observed significant upregulation of *TJP2* and *CLDN1*, core genes associated with tight junctions, following ovulatory signal stimulation, warranting further investigation into the role of tight junctions in sustaining “GC-layer stiffening”.

Focal adhesion is a protein complex that mediates cell-matrix connection, consisting of multiple components, such as talin (TLN1), vinculin (VCL), paxillin, α-actinin, focal adhesion kinase (FAK) and other proteins [22]. TLN1 and VCL, in particular, are indispensable structural proteins in focal adhesion. TLN1 activates integrins, enabling them to bind to ECM and linking them with intracellular actin. VCL interacts with TLN1 and actin, stabilizing the TLN1-actin connection. Consequently, absence either of TLN1 and VCL results in significant impairment of cell-matrix adhesion [23-25]. Therefore, we chose to knock down *TLN1* and *VCL* to disrupt focal adhesion assembly. While focal adhesion’s role in tumorigenesis is extensively studied [26], its involvement in ovulation regulation is less explored. Our findings showed that disrupting focal adhesion assembly did not did not impaired the number of ovulated oocytes (Figure 4B). However, Kitasaka *et al*. reported decreased ovulated oocytes when inhibiting FAK with Y3, suggesting a role for focal adhesion-mediated signals in ovulation [27]. We speculate that this inconsistency may be attributed to two factors. Firstly, knockdown of *TLN1* and *VCL* might have limited effects on FAK activity since FAK primarily interacts with phosphatidylinositol biphosphate and paxillin, not directly with TLN1 and VCL [28]. Secondly, despite 70% knockdown efficiency of *TLN1* and *VCL* in the follicles (Figure S1), the remaining intact focal adhesions (30%) may be sufficient to support FAK activity for ensuring a normal ovulated oocyte number. In a recent study, *GC-PXN* KO mice were generated by Vann *et al*, specifically deleting paxillin in GCs [29]. Surprisingly, these mice displayed normal estrus cycles, ovulation, and fecundity, in contrast to the luteal dysfunction observed in our *si-VCL+TLN1* mice (Figure 4). Vann *et al*. found that even in the absence of paxillin, VCL was still located at the cytomembrane, and GC proliferation, migration, and attachment were unaffected. They thus proposed that paxillin’s absence does not abrogate focal adhesion. However, we hypothesize that the divergence in phenotypes between *GC-PXN* KO and *si-VCL+TLN1* mice may be attributed to the knockout strategy employed. One possibility is that the deletion of only exons 2-5 in *GC-PXN* KO mice, despite paxillin consisting of 12 exons, leads to the production of a truncated paxillin with functionality. Another possibility is that paxillin may have lesser significance compared to VCL and TLN1 in supporting the spatial structure of focal adhesion, resulting in limited impact on focal adhesion assembly. Experimental evidence is needed to validate these hypotheses.

Overall, our study introduces the novel concept of “GC-layer stiffening” as a framework to address the fundamental question of why GCs remain trapped within the ruptured follicle after ovulation. Furthermore, it provides evidence of the evolutionary conservation of GC-layer stiffening (Figure 1G), highlighting its significance across higher mammals. These findings provide a significant advancement in our understanding of the factors that prevent GC escape from the post-ovulatory follicle.

## MATERIALS AND METHODS

### Animals

Kunming mice were purchased from the Experimental Animal Center of Huazhong Agricultural University (Wuhan, China). The mice were reared in an SPF laboratory animal house and maintained at a constant temperature of 22 ±2 °C, with 12-hour light-dark cycles (lights on from 7:00 to 19:00). They were allowed to access food and water *ad libitum*. The Hainan black goats, on the other hand, were raised at the Yazhou Beiling Black Goat Farmers’ Professional Cooperative in Sanya, China. All experiments and handling of mice and goats were conducted following the guidelines of the respective animal experimental institutions. Prior approval from the Institutional Animal Ethics Committee of Huazhong Agricultural University was obtained, with the approved protocol number being HZAUMO-2020-0103 (mouse); HZAUGO-2020-004 (goat).

### Analysis of spatial transcriptomics and RNA-seq

The spatial transcriptome data of mouse ovaries, contributed by Mantri *et al* [14], was obtained from the Gene Expression Omnibus database (Login Number: GSE240271). The bead barcode location files matched to spatial transcriptomics datasets, processed spatial metadata, and cell annotations files, were sourced from GitHub (https://github.com/madhavmantri/mouse_ovulation). The assignments of labeling the cell types and distinguishing the development stage of follicles have been completed by Mantri *et al*. We re-confirmed the identity of GCs within the pre-ovulatory follicles based on morphological information and the expression abundance of marker genes *Lhcgr* and *Adamts1*. Subsequently, transcriptome data of these GCs were extracted for bioinformatics analysis. PCA analysis was employed to assess the differences in transcription profiles. Transcriptome data for monkey and human [30, 31] were obtained from the Gene Expression Omnibus database, with login numbers GSE22776 and GSE133868, respectively. Differentially expressed genes were identified using DESeq2 software, with a significance threshold of P-value <0.05. The upregulated genes were then subjected to gene ontology (GO) analysis using the DIVAD database (USA) (https://david.ncifcrf.gov/tools.jsp), and Kyoto Encyclopedia of Genes and Genomes (KEGG) analysis using the KOBAS database (China) (http://kobas.cbi.pku.edu.cn/home.d).

### Follicle culture

Small follicles measuring 180-200 μm in diameter were isolated from ovaries using 33-gauge microneedles (KONSFI, China). The isolated follicles were cultured in 96-well plates (BKMAM, China) coated with 50 μL mineral oil (Sigma-Aldrich, USA) and placed in a 37 °C incubator with 5% CO_2_. The culture medium for maturation consisted of ɑ-MEM (Gibco, USA) supplemented with 1% ITS-G (Macklin, China), 5% FBS (Serana, Germany), 10 mIU/mL FSH (NSHF, China), and 100 U/mL penicillin/streptomycin (Servicebio, China). After 96 hours of culture, follicles reaching the pre-ovulatory stage (500-550 μm) were transferred to ovulation-inducing medium and cultured for up to 16 hours. The ovulation-inducing medium contained ɑ-MEM supplemented with 1% ITS-G, 5% FBS, 10 mIU/mL FSH, 1.5 IU/mL hCG (NSHF, China), 10 ng/mL EGF (PeproThec, USA), 5mg/mL D-Glucose (MCE, USA), and 100 U/mL penicillin/streptomycin. In experiments studying the signaling pathway, Forsklin (MCE, USA) and H-89 (MCE, USA) dissolved in DMSO were added to the ovulation induction solution at concentrations of 20 and 50 μM, respectively. After ovulation, the post-ovulatory follicles were transferred to luteal culture medium and cultured for up to 24 hours. The luteal culture medium consisted of ɑ-MEM supplemented with 1% ITS-G, 5% FBS, 10 mIU/mL FSH, 1.5 IU/mL hCG, 10 ng/mL EGF, 1ng/mL Prolactin (MCE, USA), 10 μM Cholesterol (MCE, USA), and 100 U/mL penicillin/streptomycin.

### Superovulation

To stimulate follicle growth to the pre-ovulatory stage in weaned juvenile mice, an injection of 5 IU of PMSG (Ningbo Sansheng Biological Technology, China) was administered. After 48 hours, ovulation and luteinization were triggered by injecting 5 IU of hCG (Ningbo Sansheng Biological Technology, China). In goats, vaginal plugs containing progesterone (Zoetis Australia Pty Ltd, New Zealand) were pre-inserted to synchronize their estrus cycle. For superovulation induction, the goats were injected with follicle-stimulating hormone (40 IU, Ningbo Sansheng Biological Technology, China) seven times at 12-hour intervals, starting 84 hours before the removal of the plugs. At the time of plug withdrawal, 240 IU of PMSG was injected. After an additional 14 hours, 100 IU of LH (Ningbo Sansheng Biological Technology, China) was administered, and the GCs clumps were obtained for rigidity determination by puncturing the pre-ovulatory follicles 0 and 10 hours after the LH injection.

### Determination of GC-layer rigidity

The ovaries or cultured follicles were punctured with a microneedle at the desired time point to release the GC clumps. They are then transferred to the DMEM/F12 buffer (Gibco, USA) and placed in a thermostatic shaker (Leopard, China) for 1 minute of mechanical oscillation. The oscillation parameter was set to 700 rpm at 37 °C. The oscillations were recorded and photographed using a stereo microscope (Olympus Corporation, Japan, SZX16).

### Live recording of ovulation

The pre-ovulatory follicles were cultured in a 96-well plate with 60 μl of ovulation-inducing medium per well. The plate was then placed in a 37 °C incubator with 5% CO_2_. After 8 hours of culture, the plate containing the follicles was transferred to a live-cell imaging system (Agilent BioTek Cytation 5, USA) to capture images at 6-minute intervals during ovulation. Subsequently, all the images were compiled to create a comprehensive video.

### RNA interference

Lentivirus-mediated RNA interference was used to inhibit the expression of target genes in follicles or ovaries. Briefly, PLKO.1-EGFP-PURO plasmid (Genecreate, China) was utilized to construct interference vectors. Small interfering RNAs (siRNAs) targeting *VCL*, *TLN1*, and *CREB* were synthesized by Genepharma (China), with the following targeted sequences: *VCL*-5′-ccacgatgaagctcggaaatg-3′, *TLN1*-5′-gcccattgtaatctctgctaa-3′, *CREB*-5′-cagcagctcatgcaacatcat-3′. The common negative siRNA was purchased from Sigma-Aldrich (USA). Lentiviruses were produced in 293 T cells (ATCC, USA) by co-transfecting 4.8 μg of the interference vector, 2.4 μg of pMD2.G, and 3.6 μg of pSPAX2. After 48 hours, the viral supernatants were harvested, centrifuged, and filtered through 0.45 μm polyvinylidene fluoride (PVDF) membranes (Sigma, USA). To knockdown the expression of the target genes, the follicles were cultured in maturation medium containing 10% lentivirus (titer: 1.25×10^7^ viral particles/mL) for 48 hours. For inhibiting the expression of the target gene in the ovaries, 15-day-old mice were anesthetized with 1% pentobarbital sodium. Subsequently, 2.5μl of lentivirus with a titer of 1.25 × 10^9^ viral particles/mL was injected beneath the ovarian bursa using a 10μl syringe (Hamilton, Switzerland) and a 33-gauge Small Hub RN Needle (Hamilton, Switzerland). Follow-up experiments on these mice were conducted 5 days after plasmid transfection.

### qRT-PCR analysis

Total RNA was extracted from the collected samples using the Trizol reagent. Reverse transcription was performed using the PrimeScript RT reagent kit (Takara, Japan). The qRT-PCR was conducted using a CFX384 Real-Time PCR System (Bio-Rad, USA). The reaction mixture comprised of 5 μl SYBR Green (Biosharp, China), 2 μl complementary DNA template, 250 nM each of the forward and reverse primers, and ddH_2_O to make a total volume of 10 μl. The reaction conditions were as follows: initial denaturation at 95 ℃ for 10 min, followed by 35 cycles of denaturation at 95 ℃ for 10 s, and annealing / extension at 60 ℃ for 30 s. A final step included a melting curve analysis ranging from 60 ℃ to 95 ℃, with a 0.5 ℃ increment every 5 s. Gene expression levels were normalized using the housekeeping gene *ACTB*, and the relative RNA quantification was determined using the comparative 2^−△△Ct^ method. The primer sequences used for PCR amplification are provided in Table S1.

### Frozen section and H&E staining

The ovaries or cultured corpus luteum were collected as per experimental requirements and embedded in an OCT embedding medium (Sakura, USA) for subsequent processing. The embedded tissues were flash-frozen in liquid nitrogen for 1 minute and then sectioned into 6 μm-thick slices using a frozen microtome (Leica). The sections were stained with hematoxylin and eosin (Servicebio, China) and examined under a microscope (Olympus, Japan). Photomicrographs were captured, and parameters including the number of luteal cells, the area of the corpus luteum, and the area of cavities were measured using ImageJ software. The density of luteal cells was calculated using the formula: luteal cell density = the number of luteal cells / (corpus luteum area - cavity area).

### ChIP-qPCR assay

ChIP-qPCR was employed to assess the abundance of CREB binding in the promoter regions of *VCL* and *TLN1*. The isolated GC samples were fixed in 10mL of DMEM/F-12 supplemented with 1% formaldehyde (Cell Signaling Technology, USA) for 10 minutes at room temperature with rotation. The reaction was then quenched by adding 1mL of 1.5 M glycine and rotating for an additional 5 minutes at room temperature. The samples were transferred into a 1.5 mL centrifuge tube (Axygen, USA) containing PBS for wash, and then lysed in cytomembrane lysis buffer at 4°C for 15 minutes with mixing every 5 minutes. The buffer contains 10 mM HEPES (Sigma-Aldrich) at pH 7.9, 0.5% IGEPAL-CA630 (Sigma-Aldrich, USA), 1.5 mM MgCl_2_ (Sigma-Aldrich, USA), 10 mM KCl (Sigma-Aldrich, USA), and a protease inhibitor cocktail (Sigma-Aldrich, USA). Following this, the samples were further lysed in nuclear lysis buffer, containing 1% SDS (Sigma-Aldrich, USA), 10 mM EDTA (Sigma-Aldrich, USA), 50 mM Tris at pH 8.1 (Sigma-Aldrich, USA), and a protease inhibitor cocktail, for 15 minutes at 4°C. Finally, the chromatin was sonicated using an ultrasonic disintegrator (Bioruptor PLUS, Belgium) to fragment the DNA into sizes ranging from 100 to 500 bp. The immunoprecipitation (IP) experiments were performed using the Magna ChIP™ A/G Chromatin Immunoprecipitation Kit (Merck, USA). In brief, the supernatant obtained from sonicated chromatin was diluted with ChIP IP buffer. Immunoprecipitation was performed by adding 2 mg of P-CREB antibody to protein A/G Dynabeads (Life Technologies, USA) and incubating the mixture overnight at 4°C. The antibody-bound beads were then washed, and the DNA-protein complexes were eluted and subjected to reverse crosslinking. DNA purification was carried out using the QIAquick® PCR Purification Kit (Qiagen, Germany). The amplification products were visualized by agarose gel electrophoresis (80 V, 80 mA, 75 min). The primers designed for amplifying the promoter regions of *VCL* and *TLN1* were based on the sequences of the CREB binding motifs. The specific primer sequences can be found in the table S1.

### Western Blot

Total proteins were extracted with RIPA lysis buffer (ComWin Biotech, China) supplemented with protease and phosphatase inhibitors (ComWin Biotech, China) and PMSF (Solarbio, China). The protein content was determined using the BCA Protein Assay Kit (Servicebio, China). Subsequently, the proteins were separated by polyacrylamide gel electrophoresis and transferred onto a polyvinylidene fluoride membrane. Following the transfer, the membrane was blocked with 5% skim milk powder (Nestle, Switzerland) at room temperature and then incubated overnight at 4°C with the appropriate primary antibodies, including: VCL (1:1000 dilution, Abclonal, China), TLN1 (1:1000 dilution, Abclonal, China), phosphor-PKA (1:1000 dilution, CST, USA), CREB (1:1000 dilution, Abclonal, China), phospho-CREB (1:1000 dilution, CST, USA) and α-tubulin (1:1000 dilution, Servicebio, China). The membrane was subsequently washed three times with TBST (Solarbio, China) and incubated with the appropriate HRP-conjugated secondary antibodies (goat anti-rabbit secondary antibody, 1:4000 dilution; goat anti-mouse secondary antibody, 1:4000 dilution, Biodragon-immunotech, China) for 1 hour at room temperature. After washing with TBST, the protein bands were visualized using an ECL chemiluminescent reagent kit (Servicebio, China). Images were captured using a Chemiluminescence imager (Image Quant LAS4000 mini). The protein levels were normalized to the expression of the housekeeping protein α-tubulin.

### Immunofluorescence staining

The collected GC clumps were embedded in OCT (Sakura, USA) and frozen, and then sectioned into 5 μm thick slices. The sections were rewarmed and fixed, followed by high-temperature antigen retrieval at 95-98 °C for 25 minutes using a 5% antigen retrieval buffer (Servicebio, China). Next, the sections were blocked with 10% goat serum (Boster, China) for 60 minutes at room temperature and incubated overnight at 4 °C with primary antibodies, including VCL (1:50 dilution, Abclonal, China) and TLN1 (1:50 dilution, Abclonal, China). After three washes with PBS, the sections were incubated with the appropriate fluorophore-conjugated secondary antibodies (FITC labeled goat anti-rabbit secondary antibody, 1:100 dilution; CY3-labeled goat anti-rabbit secondary antibody, 1:100 dilution, Abclonal, China) at 37 °C for 2h. Following another round of washing, the sections were imaged using a LSM800 confocal microscope system (Zeiss, Germany) and the images were processed using Zen 2.3 lite software.

### Luciferase reporter assay

To construct the reporter vectors, the promoter regions of *VCL* and *TLN1* were amplified and inserted into the PGL3-Basic luciferase reporter vector (Promega, USA) using the ClonExpress Ultra One Step Cloning Kit (Vazyme, China). Concurrently, the promoter regions of of *VCL* and *TLN1* containing single base mutations were inserted into the PGL3-Basic luciferase reporter vector using the Mut Express II Fast Mutagenesis Kit (Vazyme, China). For the construction of the *CREB* over-expression vector, the full-length coding sequence (CDS) of *CREB* was amplified and inserted into the pcDNA3.1 plasmid (Addgene, USA). HEK293T cells were seeded in a 24-well plate and incubated for 24 hours. Then, the *CREB* over-expression vector, the constructed pGL3-Basic reporter vectors, and the pRL-TK vector (Promega, USA) were co-transfected into the cells using the jetPRIME® transfection reagent (Polyplus-transfection, France) at a ratio of 96: 96: 1. After 24 hours of transfection, the cells were lysed in 100 μl lysis buffer and subjected to promoter activity assay using the dual luciferase reporter assay system (Promega, USA). The luciferase enzymatic activity was measured using a PE Enspire Multilabel Reader (PerkinElmer, USA). The primers used in this experiment are listed in Table S1.

### Hormone determination

The levels of progesterone in serum and culture medium were quantified using radioimmunoassay. Briefly, sera were obtained by centrifuging whole blood at 3000 rpm for 10 minutes and stored at −20°C. Culture medium was directly collected and stored at −20°C. Detection kits purchased from the Bioengineering Institute (Nanjing, China) were utilized for the analysis, which was conducted by the North Institute of Biological Technology (Beijing, China).

### Statistics analysis

Statistical analyses were using GraphPad Prism 10.0 (GraphPad). Data were expressed as the mean ±SD. Two-tailed unpaired Student’s t test and one-way analysis of variance followed by Tukey’s post hoc test were used to analyze the statistical significance between two groups and among multiple groups, respectively. Chi-squared test was used in the comparison between the percentages. The statistical significance was set at P-value < 0.05.

## DATA AVAILABILITY

All data are available from the corresponding author upon reasonable request.

## FUNDING

This research was supported by the Fundamental Research Funds for the Central Universities (2662023DKPY001) and the National Natural Science Foundation of China (31701301).

## SUPPORTING INFORMATION

This article contains supporting information.

## ACKNOWLEDGEMENTS

We extend our deepest appreciation and respect to Mrs. Fu Bijun, Dr. He’s mother, for her genuine care and encouragement throughout this project.

## AUTHORS’ CONTRIBUTION

C.H. conceived, designed, funded, supervised and conducted the experiments, and wrote the manuscript; X.W., J.L. and H.S. articipated in experiment design and conduction, data analysis, and involved in manuscript preparation; Y.Z., W.K. and H.W assisted with sample collection and experiments conduction; G.L. and X.L. provided advices through project implementation and improved the manuscript. All authors approved the final version.

## DECLARATION OF INTERESTS

The authors declare that they have no conflicts of interest with the contents of this article.

## Notes

### Competing Interest Statement

The authors have declared no competing interest.

## REFERENCES

1. Richard JS, Liu Z, Shimada M. Chapter 22-Ovulation. In: Plant TM, Zeleznik AJ, editors. Knobil and Neill’s Physiology of Reproduction (Fourth Edition). San Diego: Academic Press; 2015. p997–1021.

2. Robker RL, Hennebold JD, Russell DL. Coordination of ovulation and oocyte maturation: a good egg at the right time. Endocrinology. 2018;159(9):3209–3218.

3. Jaffe LA, Egbert JR. Regulation of mammalian oocyte meiosis by intercellular communication within the ovarian follicle. Annu Rev Physiol. 2017;79:237–260.

4. Zhang M, Su YQ, Sugiura K, Xia G, Eppig JJ. Granulosa cell ligand NPPC and its receptor NPR2 maintain meiotic arrest in mouse oocytes. Science. 2010;330(6002):366-369.

5. Nagyova E. The Biological Role of Hyaluronan-Rich Oocyte-Cumulus Extracellular Matrix in Female Reproduction. Int J Mol Sci. 2018;19(1):283.

6. Hernandez-Gonzalez I, Gonzalez-Robayna I, Shimada M, et al. Gene expression profiles of cumulus cell oocyte complexes during ovulation reveal cumulus cells express neuronal and immune-related genes: does this expand their role in the ovulation process?. Mol Endocrinol. 2006;20(6):1300–1321.

7. Richards JS, Liu Z, Shimada M. Immune-like mechanisms in ovulation. Trends Endocrinol Metab. 2008;19(6):191–196.

8. Richards JS, Ascoli M. Endocrine, Paracrine, and Autocrine Signaling Pathways That Regulate Ovulation. Trends Endocrinol Metab. 2018;29(5):313–325.

9. Jeppesen JV, Kristensen SG, Nielsen ME, et al. LH-receptor gene expression in human granulosa and cumulus cells from antral and preovulatory follicles. J Clin Endocrinol Metab. 2012;97(8):E1524–E1531.

10. Dos Santos EC, Lalonde-Larue A, Antoniazzi AQ, et al. YAP signaling in preovulatory granulosa cells is critical for the functioning of the EGF network during ovulation. Mol Cell Endocrinol. 2022;541:111524.

11. Fan HY, Liu Z, Shimada M, et al. MAPK3/1 (ERK1/2) in ovarian granulosa cells are essential for female fertility. Science. 2009;324(5929):938–941.

12. Diaz FJ, Wigglesworth K, Eppig JJ. Oocytes are required for the preantral granulosa cell to cumulus cell transition in mice. Dev Biol. 2007;305(1):300–311.

13. Wang X, Zhou S, Wu Z, et al. The FSH-mTOR-CNP signaling axis initiates follicular antrum formation by regulating tight junction, ion pumps, and aquaporins. J Biol Chem. 2023;299(8):105015.

14. Mantri M, Zhang HH, Spanos E, Ren YA, De Vlaminck I. A spatiotemporal molecular atlas of the ovulating mouse ovary. Proc Natl Acad Sci U S A. 2024;121(5):e2317418121.

15. Zaniker EJ, Babayev E, Duncan FE. Common mechanisms of physiological and pathological rupture events in biology: novel insights into mammalian ovulation and beyond. Biol Rev Camb Philos Soc. 2023;98(5):1648–1667.

16. Chaudhuri O, Cooper-White J, Janmey PA, Mooney DJ, Shenoy VB. Effects of extracellular matrix viscoelasticity on cellular behaviour. Nature. 2020;584(7822):535-546.

17. Saraswathibhatla A, Indana D, Chaudhuri O. Cell-extracellular matrix mechanotransduction in 3D. Nat Rev Mol Cell Biol. 2023;24(7):495–516.

18. Wang X, Ji L, Wang J, Liu C. Matrix stiffness regulates osteoclast fate through integrin-dependent mechanotransduction. Bioact Mater. 2023;27:138–153.

19. Paluch EK, Aspalter IM, Sixt M. Focal adhesion-independent cell migration. Annu Rev Cell Dev Biol. 2016;32:469–490.

20. Resnik N, Sepcic K, Plemenitas A, Windoffer R, Leube R, Veranic P. Desmosome assembly and cell-cell adhesion are membrane raft-dependent processes. J Biol Chem. 2011;286(2):1499–1507.

21. Fontao L, Stutzmann J, Gendry P, Launay JF. Regulation of the type II hemidesmosomal plaque assembly in intestinal epithelial cells. Exp Cell Res. 1999;250(2):298–312.

22. Wehrle-Haller B. Structure and function of focal adhesions. Curr Opin Cell Biol. 2012;24(1):116–124.

23. Zhao Y, Lykov N, Tzeng C. Talin-1 interaction network in cellular mechanotransduction (Review). Int J Mol Med. 2022;49(5):60.

24. Sadeghian F, Ibrahim I, Ravichandran L, et al. An integrin binding motif in TLN-1/talin plays a minor role in motility and ovulation. MicroPubl Biol. 2023;2023:10.17912

25. Bays JL, DeMali KA. Vinculin in cell-cell and cell-matrix adhesions. Cell Mol Life Sci. 2017;74(16):2999–3009.

26. Zhang Y, Liu S, Zhou S, et al. Focal adhesion kinase: Insight into its roles and therapeutic potential in oesophageal cancer. Cancer Lett. 2021;496:93–103.

27. Kitasaka H, Kawai T, Hoque SAM, Umehara T, Fujita Y, Shimada M. Inductions of granulosa cell luteinization and cumulus expansion are dependent on the fibronectin-integrin pathway during ovulation process in mice. PLoS One. 2018;13(2):e0192458.

28. Tapial Martínez P, López Navajas P, Lietha D. FAK structure and regulation by membrane interactions and force in focal adhesions. Biomolecules. 2020;10(2):179.

29. Vann K, Weidner AE, Walczyk AC, Astapova O. Paxillin knockout in mouse granulosa cells increases fecundity. Biol Reprod. 2023;109(5):669–683.

30. Xu F, Stouffer RL, Müller J, et al. Dynamics of the transcriptome in the primate ovulatory follicle. Mol Hum Reprod. 2011;17(3):152–165.

31. Poulsen LC, Bøtkjær JA, Østrup O, et al. Two waves of transcriptomic changes in periovulatory human granulosa cells. Hum Reprod. 2020;35(5):1230–1245.

