## Supplementary data for "Granulosa cell-layer stiffening prevents the granulosa cells from escaping the post-ovulatory follicle"

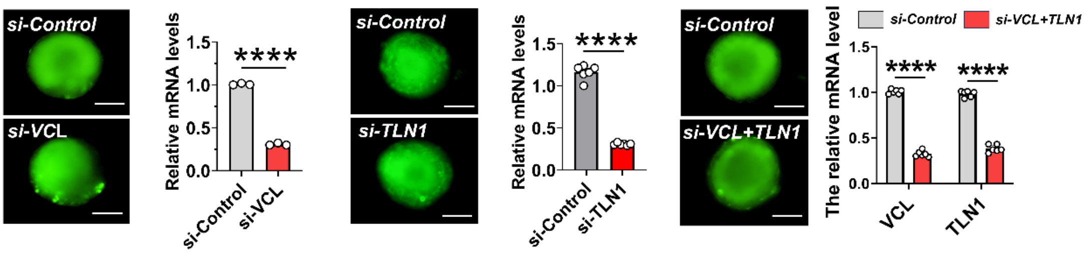

**Figure S1. qRT-PCR analysis of the efficiency of RNA interference (related to Figure 3).** Green fluorescence indicated successful transcription of interfering plasmids in follicles. Scale bar: 200 μm, n = 3 (*si-VCL*), 6 (*si-TLN1*), 6 (*si-VCL+TLN1*) follicular samples. Significance was determined using two-tailed unpaired Student's t-test, values were mean ± SD. Significant differences were denoted by ****P<0.001. The experiments were repeated three times, and similar results were obtained.

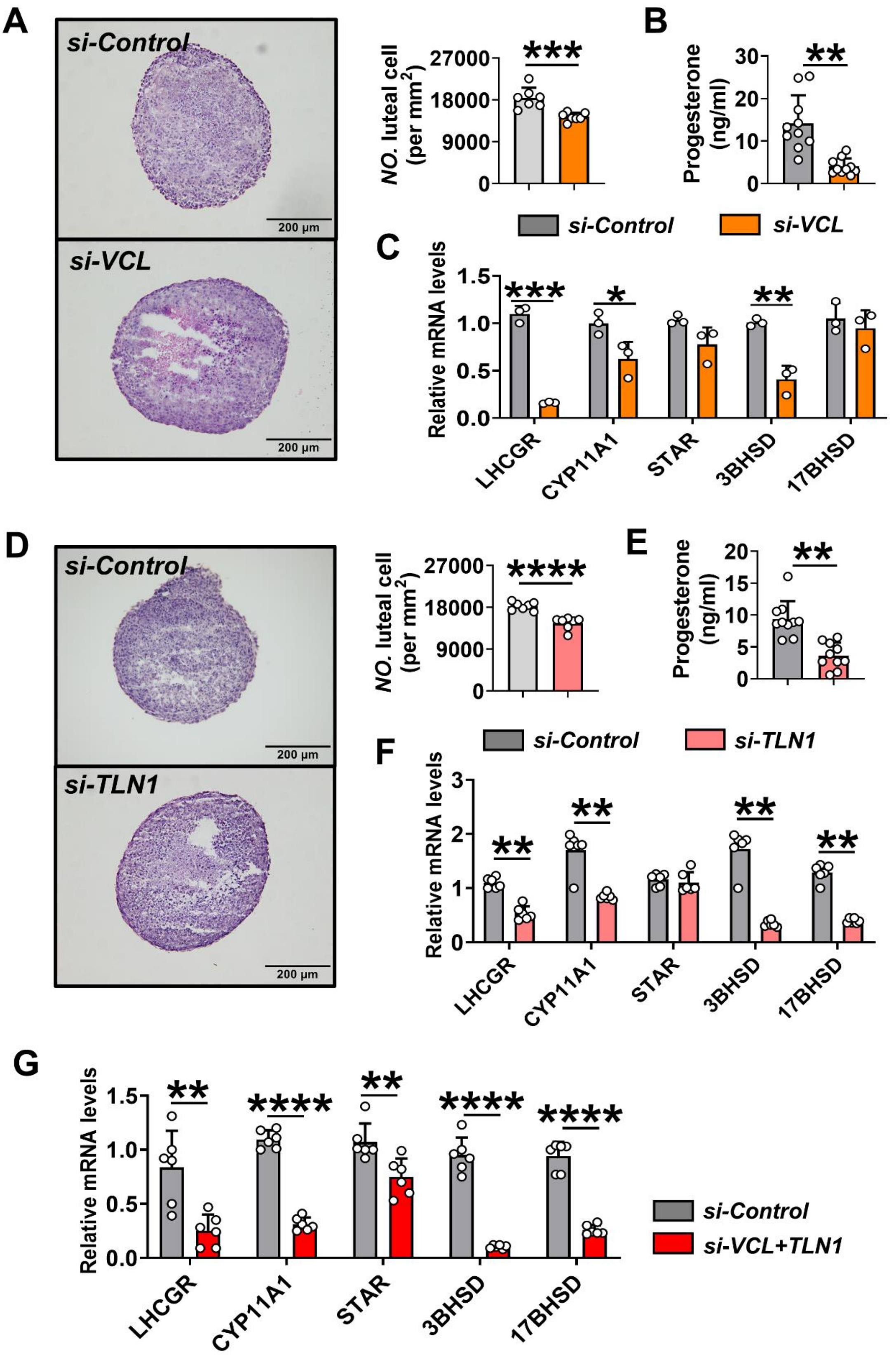

**Figure S2. Effect of *VCL* and *TLN1* knockdown on the morphology and function of culture corpus luteum (related to Figure 4).** A, Effect of *VCL* knockdown on the luteal cell density. *Left:* representative photographs of corpus luteum sections in each group, scale bar: 200 µm; *right*:

statistics of the density of luteum cells in each group (n= 7). B, Effect of *VCL* knockdown on progesterone level in culture medium (n=10). C, Change in the expression of luteal functional genes after *VCL* knockdown (n=3). D, Effect of *TLN1* knockdown on the luteal cell density. *Left:* representative photographs of corpus luteum sections in each group, scale bar: 200 µm; *right:* statistics of the density of luteum cells in each group (n= 7). E, Effect of *TLN1* knockdown on progesterone level in culture medium (n=10). F, Change in the expression of luteal functional genes after *TLN1* knockdown (n=6). G, Change in the expression of luteal functional genes after *VCL*+*TLN1* knockdown (n=6). Significance was determined using two-tailed unpaired Student's t-test, values were mean ± SD. Significant differences were denoted by *P<0.05, **P<0.01, ***P<0.001, ****P<0.001. The experiments were repeated three times, and similar results were obtained.

A

| Motif | VCL promoter SCORE | TLN1 promoter SCORE | ACTN1 promoter SCORE | ACTN4 promoter SCORE | ITGA2 promoter SCORE | ITGB1 promoter SCORE |
|---|---|---|---|---|---|---|
| CREB | 9.71 | 9.02 | 8.96 | 11.78 | 9.77 | 9.63 |
| NFKB1 | No date | No date | 11.49 | No date | 11.38 | No date |
| PPARA | 12.91 | No date | No date | No date | 12.91 | No date |
| NFATC1 | No date | 11.67 | No date | 10.79 | No date | 13.77 |
| MYC | No date | 12.65 | 10.71 | No date | No date | No date |
| FOS | 9.17 | 9.68 | No date | 9.35 | 10.66 | 8.68 |

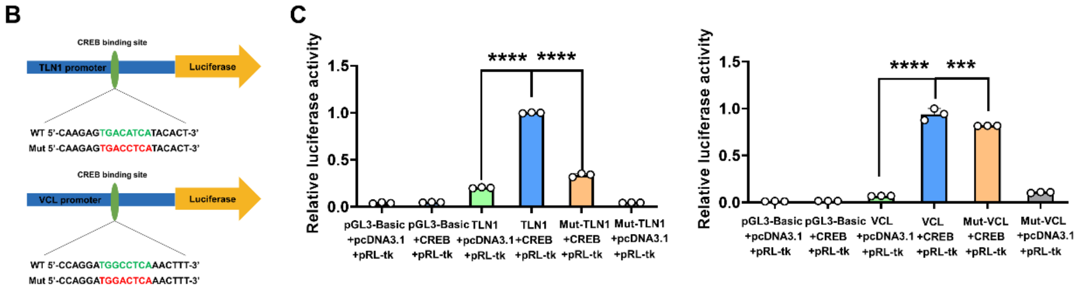

**Figure S3. *cAMP-PKA* activation and inhibition experiment and dual-luciferase experiments (related to Figure 5).** A, Binding motifs of CREB, NFKB1, PPARA, NFATC1, MYC, FOS from JASPAR database were detected in the promoter region of *VCL*, *TLN1*, *ACTN1*, *ACTN4*, *ITGA2*, *ITGB1*. B, Luciferase reporters of WT VCL/TLN1and Mut VCL/TLN1 were constructed. C, WT or Mut VCL, TLN1 Promoter Luc was measured in HEK293T cells with *CREB* overexpression. and the empty pcDNA3.1 vector was used as a negative control in Dual-Luciferase experiments. (n=3). Statistical significance was determined using two-tailed unpaired Student's t test, values were mean ± SD. Significant differences were denoted by ***P<0.001, ****P<0.0001. The experiments of C were repeated three times, and similar results were obtained.

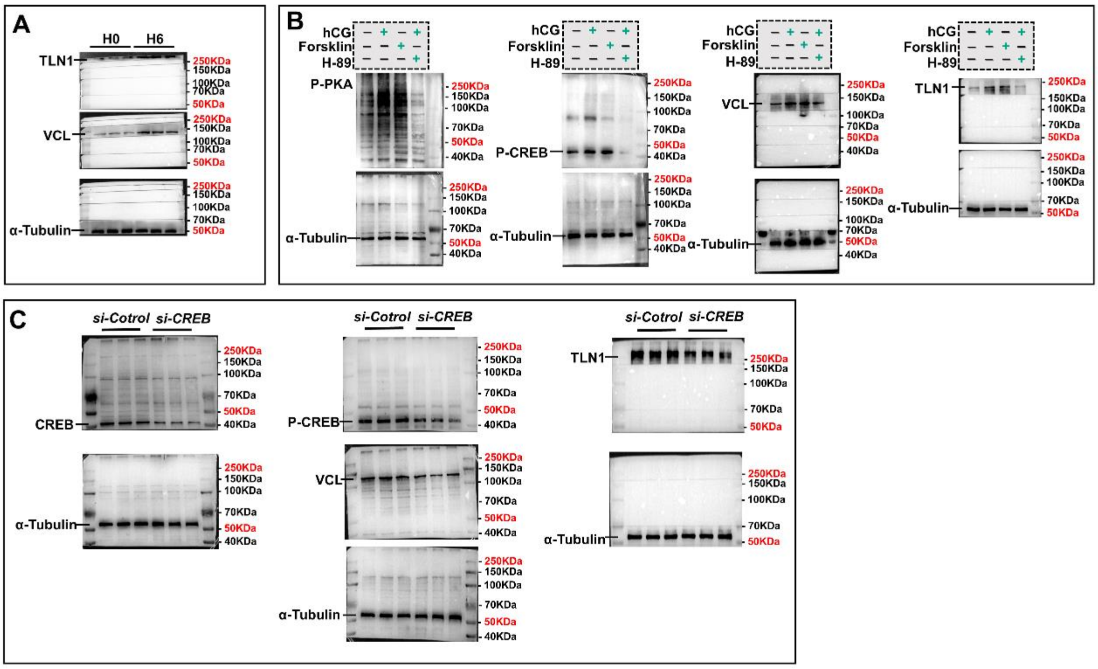

**Figure S4. The full western blots.** A, Western blot assay of the protein contents of VCL and TLN1 after hCG injection (related to Figure 2H). B, Western blot assay of the protein contents of VCL and TLN1 following activation or inhibition of the *cAMP-PKA* cascade (related to Figure 5C). C, western blot assay of protein contents of VCL and TLN1 after CREB knockdown (related to Figure 5F).

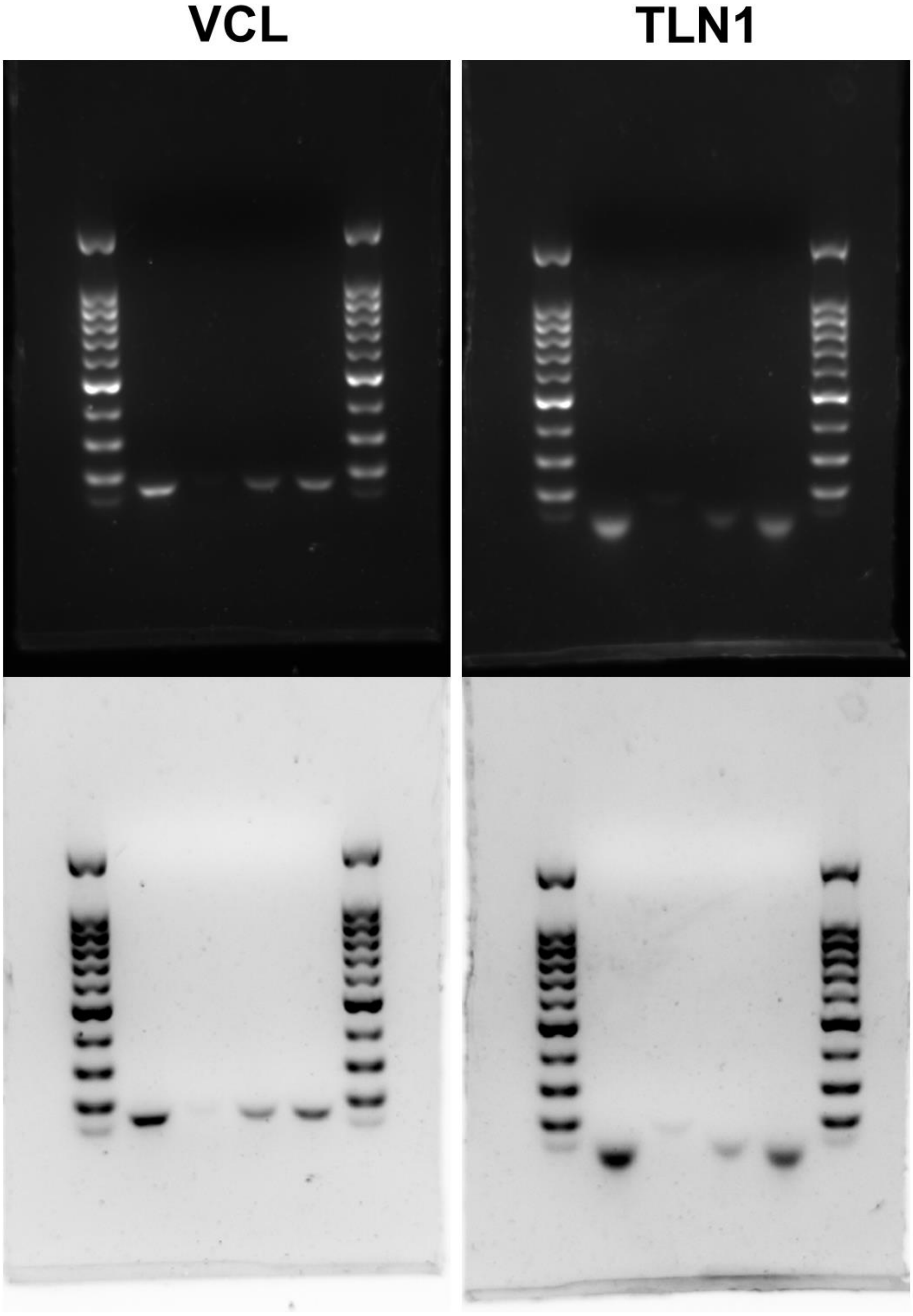

**Figure S5. Original agarose gel eletrophoresis in Figure 5E.**

**Table S1. The primers used for qPCR, ChIP-qPCR and Luciferase reporter**

| Gene | Primer sequence (5'-3') |
|---|---|
| *Tjp1-mouce* | Forward: GCCGCTAAGAGCACAGCAA<br>Reverse: TCCCCACTCTGAAAATGAGGA |
| *Tjp2-mouce* | Forward: ATGGGAGCAGTACACCGTGA<br>Reverse: TGACCACCCTGTCATTTTCTTG |
| *Tjp3-mouce* | Forward: TCGGCATAGCTGTCTCTGGA<br>Reverse: GTTGGCTGTTTTGGTGCAGG |
| *Cldn1-mouce* | Forward: GGGGACAACATCGTGACCG<br>Reverse: AGGAGTCGAAGACTTTGCACT |
| *Cldn2-mouce* | Forward: CAACTGGTGGGCTACATCCTA<br>Reverse: CCCTTGGAAAAGCCAACCG |
| *Cldn3-mouce* | Forward: ACCAACTGCGTACAAGACGAG<br>Reverse: CAGAGCCGCCAACAGGAAA |
| *Cldn4-mouce* | Forward: GTCCTGGGAATCTCCTTGGC<br>Reverse: TCTGTGCCGTGACGATGTTG |
| *Cldn6-mouce* | Forward: ATGGCCTCTACTGGTCTGCAA<br>Reverse: GCCAACAGTGAGTCATACACCTT |
| *Ocln-mouce* | Forward: TTGAAAGTCCACCTCCTTACAGA<br>Reverse: CCGGATAAAAAGAGTACGCTGG |
| *Jup-mouce* | Forward: TGGCAACAGACATACACCTACG<br>Reverse: GGTGGTAGTCTTCTTGAGTGTG |
| *Dsp-mouce* | Forward: GGATTCTTCTAGGGAGACTCAGT<br>Reverse: TCCACTCGTATTCCGTCTGGG |
| *Dsg1a-mouce* | Forward: ACTGTGTTAAATGTCATCGAGGG<br>Reverse: TGCCTGTTCTTGAGTCAACAAC |
| *Dsg1b-mouce* | Forward: GCAGTGGTGGTAATCGTGACC<br>Reverse: GGATTTTGCCTACCGGGAGTG |
| *Dsg2-mouce* | Forward: GTGGTCTGCTTGGACTTTGGA<br>Reverse: GGAACGGTTTGCCTTCATTTC |
| *Dsg3-mouce* | Forward: TGGCAGTCTGGAAGTCACC<br>Reverse: CTGTAGAGGGTCAGGGATGG |
| *Dsc1-mouce* | Forward: GGTCAAGGAATCAAAACACAGC<br>Reverse: CCAAGCCGAGGTTGAGTGAAA |
| *Dsc2-mouce* | Forward: ATGGCGGCTGTGGGATCTAT<br>Reverse: GCAAGGATCGCAAGGGTCAA |
| *Dsc3-mouce* | Forward: AGTTTGAAAGAGTGTCTCAGCTC<br>Reverse: ACAACAGCTCTGGTCGGATAA |
| *Vcl-mouce* | Forward: TGGACGGCAAAGCCATTCC<br>Reverse: GCTGGTGGCATATCTCTCTTCAG |
| *Tln1-mouce* | Forward: CCTGCCGCATGATTCGTGA<br>Reverse: TCGGAGCATGTAGTAGTCCAAA |
| *Actn1-mouce* | Forward: GACCATTATGATTCCCAGCAGAC<br>Reverse: CGGAAGTCCTCTTCGATGTTCTC |

| | |
|---|---|
| *Actn4-mouce* | Forward: ATGGTGGACTACCACGCAG<br>Reverse: CAGCCTTCCGAAGATGAGAGT |
| *Ptk2-mouce* | Forward: GAGTACGTCCCTATGGTGAAGG<br>Reverse: CTCGATCTCTCGATGAGTGCT |
| *Ptk2b-mouce* | Forward: TGAGCCCTTGAGCCGTGTA<br>Reverse: AGCTTGAAGTTCTTCCCTGGG |
| *Pxn-mouce* | Forward: CAAACGGCCAGTGTTCTTGTC<br>Reverse: TGTGTGGTTTCCAGTTGGGTA |
| *Itga2-mouce* | Forward: TGTCTGGCGTATAATGTTGGC<br>Reverse: CTTGTGGGTTCGTAAGCTGCT |
| *Itgb1-mouce* | Forward: ATGCCAAATCTTGCGGAGAAT<br>Reverse: TTTGCTGCGATTGGTGACATT |
| *Star-mouce* | Forward: GTGAAGGCTAAGGGATAA<br>Reverse: TGGAGCTGGTAAGACAAC |
| *17β-Hsd-mouce* | Forward: CCACCTGTGTTTGGCGTGTA<br>Reverse: GAGGTTGAATTGTGGATTAGGCA |
| *Cyp11a1-mouce* | Forward: GGGCAGTTTGGAGTCAGTTTAC<br>Reverse: TTTAGGACGATTCGGTCTTTCTT |
| *3β-Hsd-mouce* | Forward: TGGACAAAGTATTCCGACCAGA<br>Reverse: GGCACACTTGCTTGAACACAG |
| *Lhcgr-mouce* | Forward: CTGAGGAGATTTGGTTGCTGTA<br>Reverse: ATTTGGGTGGACTTTTTTGGGG |
| *β-Actin-mouce* | Forward: CCAGCCTTCCTTCTTGGGTAT<br>Reverse: AGGTCTTTACGGATGTCAACG |
| *Gapdh-mouce* | Forward: AGGTCGGTGTGAACGGATTTG<br>Reverse: TGTAGACCATGTAGTTGAGGTCA |
| *Vcl-goat* | Forward: GGGTCTTGGAAGCCTAGTGG<br>Reverse: AGGCAGACTGGGTTTCTAGC |
| *Tln1-goat* | Forward: GACTGAGGTACAGCAGCGTT<br>Reverse: CGGACTCCAAGTGCCTTCAT |
| *Actn1-goat* | Forward: GGGCTTGGTTTCACGCTCTG<br>Reverse: GTAAACTGTCACTTCACGGGCA |
| *β-Actin-goat* | Forward: CCTGCGGCATTCACGAAACTAC<br>Reverse: ACAGCACCCTGTTGGCGTAGAG |
| *Gapdh-goat* | Forward: GCAAGTTCCACGGCACAG<br>Reverse: GGTTCACGCCCATCACAA |
| *VCL-CREBmotif* | Forward: GCCTCTGGAGTGCTGTTGTA<br>Reverse: AGGGAAGTGGATCAGGGGTT |
| *TLN1-CREBmotif* | Forward: CTCTGATAATGCCTGTCAAACTTGG<br>Reverse: TGCTTCTAACTCCCTCGGACT |
| *INFU-VCL* | Forward: gagctcttacgcgtgctagcCTACCTGCTTCTGTCTCTGAAGTGC<br>Reverse: acagtaccggaatgccaagcCAGCCCAGAGAAATCGGCA |
| *INFU-TLN1* | Forward: gagctcttacgcgtgctagcATACTCTCCACAGCCTCTCTGTCAG<br>Reverse: acagtaccggaatgccaagcCCGAGTTGGAGAGAACTCCA |

| | |
|---|---|
| *Mut-INFU-VCL* | Forward: CAGGATGGaCTCAAACTTTCTTTGTAGTGGAGTATGA |
| | Reverse: AGTTTGAGtCCATCCTGGAGGGGTAGTCCTTT |
| *Mut-INFU-TLN1* | Forward: AGAGTGACcTCATACACTTGTAATCTAGTCCGAGGG |
| | Reverse: GTGTATGAgGTCACTCTTGGCTAGTGTCCAAGTT |
